# Oxford Nanopore Sequencing, Hybrid Error Correction, and *de novo* Assembly of a Eukaryotic Genome

**DOI:** 10.1101/013490

**Authors:** Sara Goodwin, James Gurtowski, Scott Ethe-Sayers, Panchajanya Deshpande, Michael C. Schatz, W. Richard McCombie

**Affiliations:** Cold Spring Harbor Laboratory, One Bungtown Road. Cold Spring Harbor, NY 11724

**Keywords:** Single molecule sequencing, Oxford Nanopore, Genome assembly, yeast

## Abstract

Monitoring the progress of DNA molecules through a membrane pore has been postulated as a method for sequencing DNA for several decades. Recently, a nanopore-based sequencing instrument, the Oxford Nanopore MinION, has become available that we used for sequencing the *S. cerevisiae* genome. To make use of these data, we developed a novel open-source hybrid error correction algorithm Nanocorr (https://github.com/jgurtowski/nanocorr) specifically for Oxford Nanopore reads, as existing packages were incapable of assembling the long read lengths (5-50kbp) at such high error rate (between ∼5 and 40% error). With this new method we were able to perform a hybrid error correction of the nanopore reads using complementary MiSeq data and produce a *de novo* assembly that is highly contiguous and accurate: the contig N50 length is more than ten-times greater than an Illumina-only assembly (678kb versus 59.9kbp), and has greater than 99.88% consensus identity when compared to the reference. Furthermore, the assembly with the long nanopore reads presents a much more complete representation of the features of the genome and correctly assembles gene cassettes, rRNAs, transposable elements, and other genomic features that were almost entirely absent in the Illumina-only assembly.

**Reviewer link to data:** http://schatzlab.cshl.edu/data/nanocorr/

## Introduction

Most DNA sequencing methods are based on either chemical cleavage of DNA molecules^1^, or synthesis of new DNA strands^2^, which is used in the majority of today’s sequencing routines. In the more common synthesis based methods, base analogues of one form or another are incorporated into a nascent DNA strand that is labeled either on the primer from which it originates or on the newly incorporated bases. This is the basis of the sequencing method used for most current sequencers, including Illumina, Ion Torrent, and PacBio sequencing, and their earlier predecessors^3^. Alternatively, it is been observed that individual DNA molecules could be sequenced by monitoring their progress through various types of pores^4,5^, originally envisioned as being pores derived from bacteriophage particles^6^. The advantages of this approach include potentially very long and unbiased sequence reads as no amplification nor chemical reactions are necessary for sequencing^7^.

Recently we began testing a sequencing device using nanopore technology from Oxford Nanopore Technologies through their early access program^8^. This device, the MinION, is a nanopore-based device in which pores are embedded in a membrane placed over an electrical detection grid. As DNA molecules pass through the pores they create measureable alterations in the ionic current. The fluctuations are sequence dependent and thus can be used by a base-calling algorithm to infer the sequence of nucleotides in each molecule^7,9^. As part of the library preparation protocol, a hairpin adapter is ligated to one end of a double stranded DNA sample while a “motor” protein is bound to the other to unwind the DNA and control the rate of nucleotides passing through the pore^10^. Under ideal conditions the leading template strand passes through the pore followed by the hairpin adapter and then the complement strand. In such a run where both strands are sequenced, a consensus sequence of the molecule can be produced; these consensus reads are termed “2D reads” and, have generally higher accuracy than reads from only a single pass of the molecule (“1D reads”)^11^.

The ability to generate very long read lengths from a handheld sequencer opens the potential for many important applications in genomics, including *de novo* genome assembly of novel genomes, structural variation analysis of healthy or diseased samples, or even isoform resolution when applied to cDNA sequencing. However, both the 1D and 2D read types currently have a high error rate that limits their direct application to these problems, and necessitate a new suite of algorithms. Here we report our experiences sequencing the *S. cerevisiae* (yeast) genome with the instrument, including an in-depth analysis of the data characteristics and error model. We also describe our new hybrid error correction algorithm Nanocorr, that leverages high quality short read MiSeq sequencing to computational “polish” the long Nanopore reads. After error correction, we then *de novo* assemble the genome using just the error corrected long reads to produce a very high quality assembly of the genome with each chromosome assembled into a small number of contigs at very high sequence identity. We further demonstrate our error correction is nearly optimal: our results with the error corrected real data approach those produced using idealized simulated reads extracted directly from reference genome itself. Finally, we validate these results by error correcting long Oxford Nanopore reads of the E. coli K12 genome sequenced at a different institution and produce an essentially perfect *de novo* assembly of the genome. As such, we believe our hybrid error correction and assembly approach will be generally applicable to many other sequencing projects.

## Results

### Nanopore Sequencing of Yeast

We chose to sequence the yeast genome with the new nanopore sequencer so that we could carefully measure the accuracy and other data characteristics of the device on a tractable and well-understood genome. Our initial flow cells had somewhat low reliability and throughput, but improved substantially over time (**Supplemental Figure S1**). This is due to a combination of improvements in chemistry, protocols, instrument software, and shipping conditions. Some runs have produced upwards of 450 Mb of sequencing data per flow cell over a 48 hour period. All together, we generated more than 195x coverage of the genome with an average read length of 5,548bp but with a long tail extending to a maximum read length of 191,145bp for a “1D read” and 57,453bp for a “2D read” (**Supplemental Note 3**). These reads derived from three separate iteration of the device: R6.0, the earliest version of the device, accounts for ∼11% of the data produced in this study; the R7.0 iteration of the device accounts for ∼49% of the data; and R7.3, the most recent version of this device, accounts for ∼40% of the data produced.

Alignment of the reads to the reference genome using BLAST gave us a deeper analysis of the per base error rate. Of the 361,647 reads produced by our 46 sequencing runs, 44,028 2D reads (or about 56% of the 2D reads) and 105,771 1D reads (about 31% of the 1D reads) aligned to the reference yeast genome. The remaining reads either mapped to a control sequence used as a spike-in for some experiments (about 8.5% of the reads) or did not show significant similarity to the W303 genome or spike-in sequence, presumably because of insufficient read quality (**Supplemental Note 4**). The mean identity to the reference of “1D” reads was between 58.8% (R6.0 flowcells) and 64.60% (R7.3 flowcells) while the average “2D” read identity was between 60.96% (R6.0) and 75.39% (R7.3) with many 2D reads exceeding 80% identity **(Supplemental Figure S5A)**. The overall alignment identities of both 1D and 2D reads are summarized in **Figure 1A** that compares both read length and percent identity. Other aligners, including LAST, were also tested and gave comparable results (**Supplemental Note 9**).

**Figure 1.**
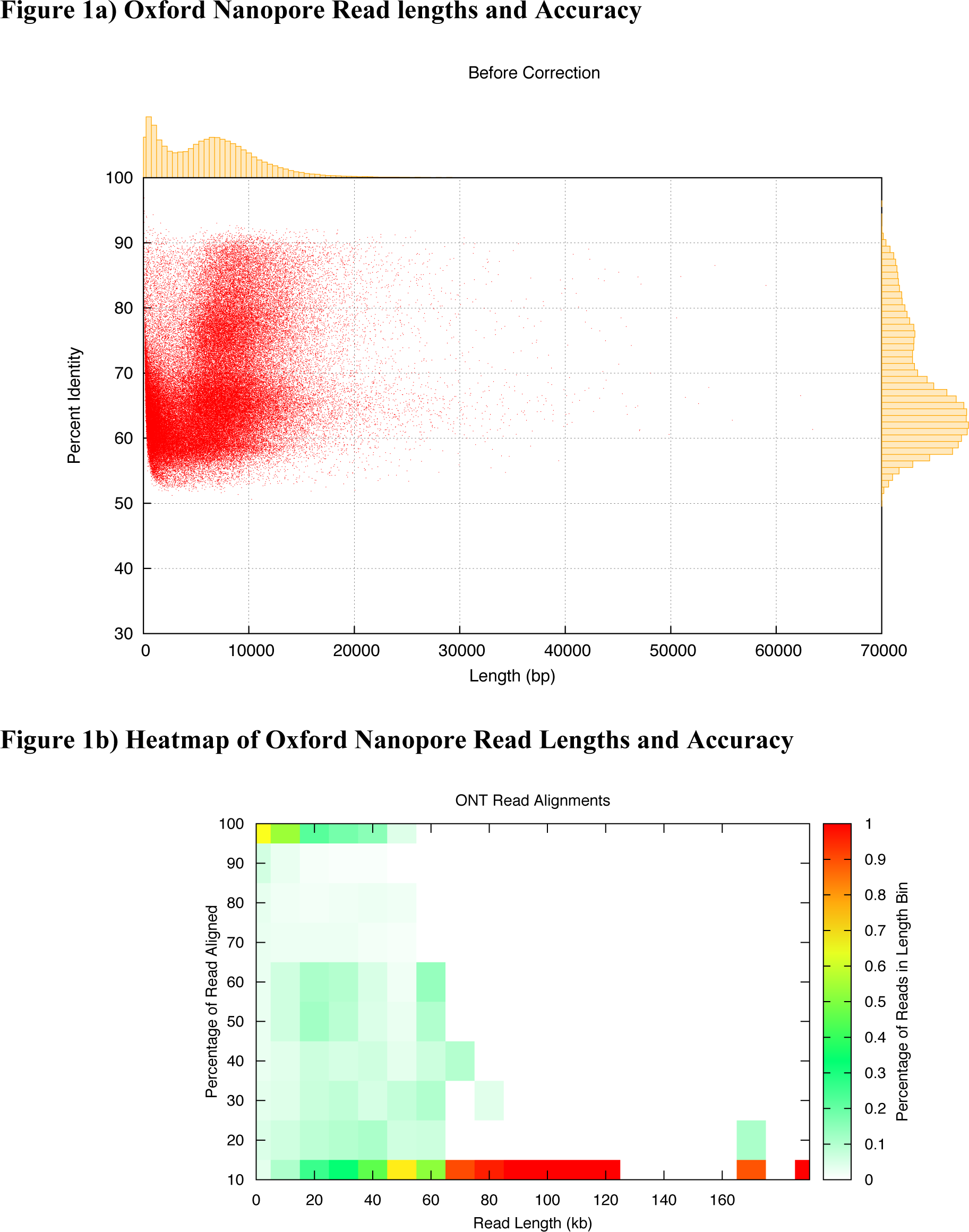
A) Scatter plot of read length versus accuracy with marginal histograms summarizing the raw ONT alignments. B) Heatmap of read length versus percent of read aligned to the S288C genome. Each cell represents a summary of how reads of different lengths align. Each color represents the fraction of reads in a given read length bin. Maximal alignment efficiency is observed between 10 and about 40 kb, while fragments longer than 80kb are virtually unalignable.

Overall read quality is further summarized by **Figure 1B** showing a heatmap of the lengths of the alignments relative to the full length of the reads. On the lower end (below 50kbp), a substantial number (up to 50%) of the reads do not align to the reference in any capacity. However, those that can be aligned have matches that span nearly their entire length. For longer reads (>50kb), only portions of the reads can be successfully aligned which suggests that reads are composed of both high and low quality segments. However, this local variability in quality does not seem to be position specific, and on average the per-base error rate is consistent across the length of a read **(Supplemental Figure S5B)**. The very longest reads tend not to be alignable at all, suggesting that the longest reads may be extremely low quality or include other artifacts of the sequencing process.

Evaluating a sample of the aligned reads, the overall coverage distribution approximated a Poisson distribution, although some over-dispersion was observed that was better modeled by a Negative Binomial distribution **(Supplemental Figure S5C)**. To examine some of the sources of the over-dispersion we also examined the coverage as a function of the GC composition of the genome. Between 20% and 60% GC content, the coverage was essentially uniform, while at higher and lower GC content the coverage is more variable partially explaining some of the regions of the genome lacking raw read coverage **(Supplemental Figure S5D)**.

### Hybrid Error Correction and de novo Assembly

To demonstrate the utility of the long reads, we attempted to assemble the yeast genome *de novo* using the Celera Assembler, which can assemble low-error rate reads up to 500kbp long. However, when raw nanopore reads were given to the assembler, not one single contig was assembled, and it became apparent that error correction was critical to the success of the assembly. Consequently, we developed a novel algorithm called Nanocorr to error correct the reads prior to *de novo* assembly or other purposes. Nanocorr uses a hybrid strategy for error correction, using high quality MiSeq short reads to error correct the long but highly erroneous nanopore reads. It follows the design of hybrid error correction pipelines for PacBio long read sequencing^12^, although in our testing none of the available algorithms were capable of utilizing the nanopore reads. For example, the HGAP error correction algorithm for PacBio reads produced 2318 reads (0.18x coverage) while the hybrid PacBio/Illumina error correction algorithm PacbioToCA produced only 167 reads (0.06x). We were therefore motivated to develop an entirely new algorithm.

Briefly, Nanocorr begins by aligning the short MiSeq reads to the long nanopore reads using the BLAST sequence aligner. This produced a mix of correct, near-full length alignments along with false or partial alignments of the short reads. To separate these types of alignments, Nanocorr uses a dynamic programming algorithm based on the longest-increasing-subsequence (LIS) problem to select the optimal set of short read alignments that span each long read. The consensus reads are then calculated using a finite state machine of the most commonly observed sequence transitions using the open source algorithm *pbdagcon*^13^ **(Figure 3A)**. Overall, we find this process increased the percent identity from an average of 67% for uncorrected reads from flowcell iterations R6.0-R7.3 to over 97% (**Figure 2B**, **Supplemental Figure S6A**). The error corrected long reads can be used for any purpose, especially *de novo* genome assembly.

**Figure 2.**
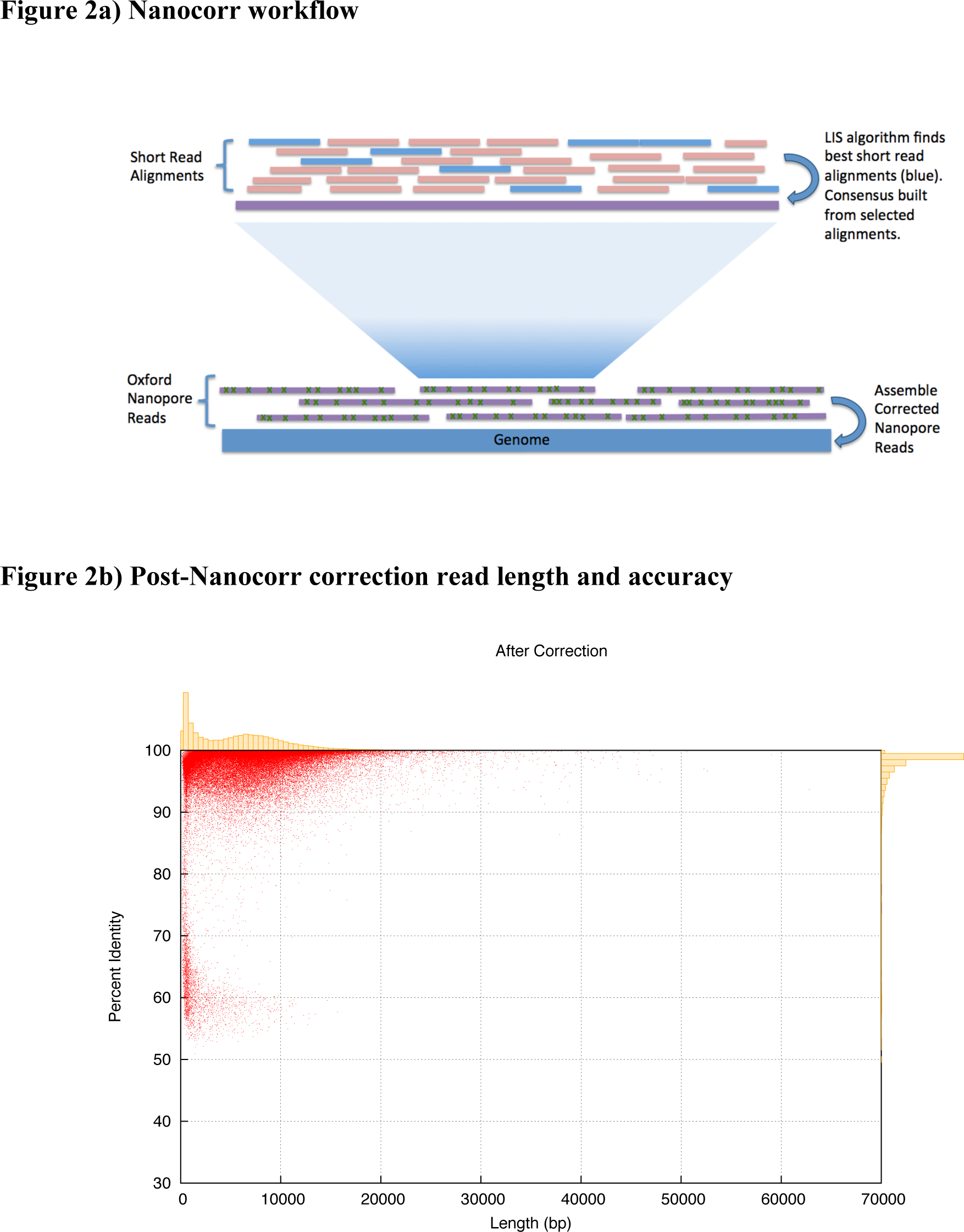
A) Nanocorr workflow. Short high identity reads are aligned to raw ONT reads. The best overlapping set is determined by the LIS algorithm and a consensus sequence of these alignments is built using pbdagcon. Error corrected reads can then be assembled using a long read assembler. B) Scatterplot with marginal histograms summarizing the percent identity of reads after correction for W303. Average identity before correction is ∼68% for all iterations of flowcells, while the average post-correction identity was over 97%.

After error correction, we selected the set of reads that were greater than 4kb in length from the three highest yielding flowcells (**Supplemental Note 10**). This brought us to our target of ∼20x coverage of the genome, for *de novo* assembly. We then assembled those reads with the Celera Assembler, which follows an overlap-layout-consensus approach without decomposing the long reads into *k-*mers as is used in de Bruijn graph assemblers. This produced an assembly consisting of 108 non-redundant contigs with a N50 size of 678kbp and requiring only a few contigs needed to span each chromosome (**Supplemental Figure F6B**). Upon alignment to the reference sequence we found that over 99% of the reference genome aligned to our assembly and the per-base accuracy of our assembly was more than 99.78%. Furthermore, after polishing the assembly with the algorithm Pilon^14^ the per-base identity was further improved to 99.88%. We investigated the residual differences, and found the majority of differences between the nanopore assembly and the S288C reference genome reside in repetitive regions, especially long repetitive regions and homopolymer sequences, while the accuracy of gene sequences was over 99.9%. The assembly also has substantially better resolution of the genome compared to an assembly of the MiSeq reads on their own which has a contig N50 size of only 59kbp: the Nanopore-based assembly is more than an order of magnitude more contiguous across all cutoffs in the contig length distribution (**Figure 3**)

**Figure 3.**
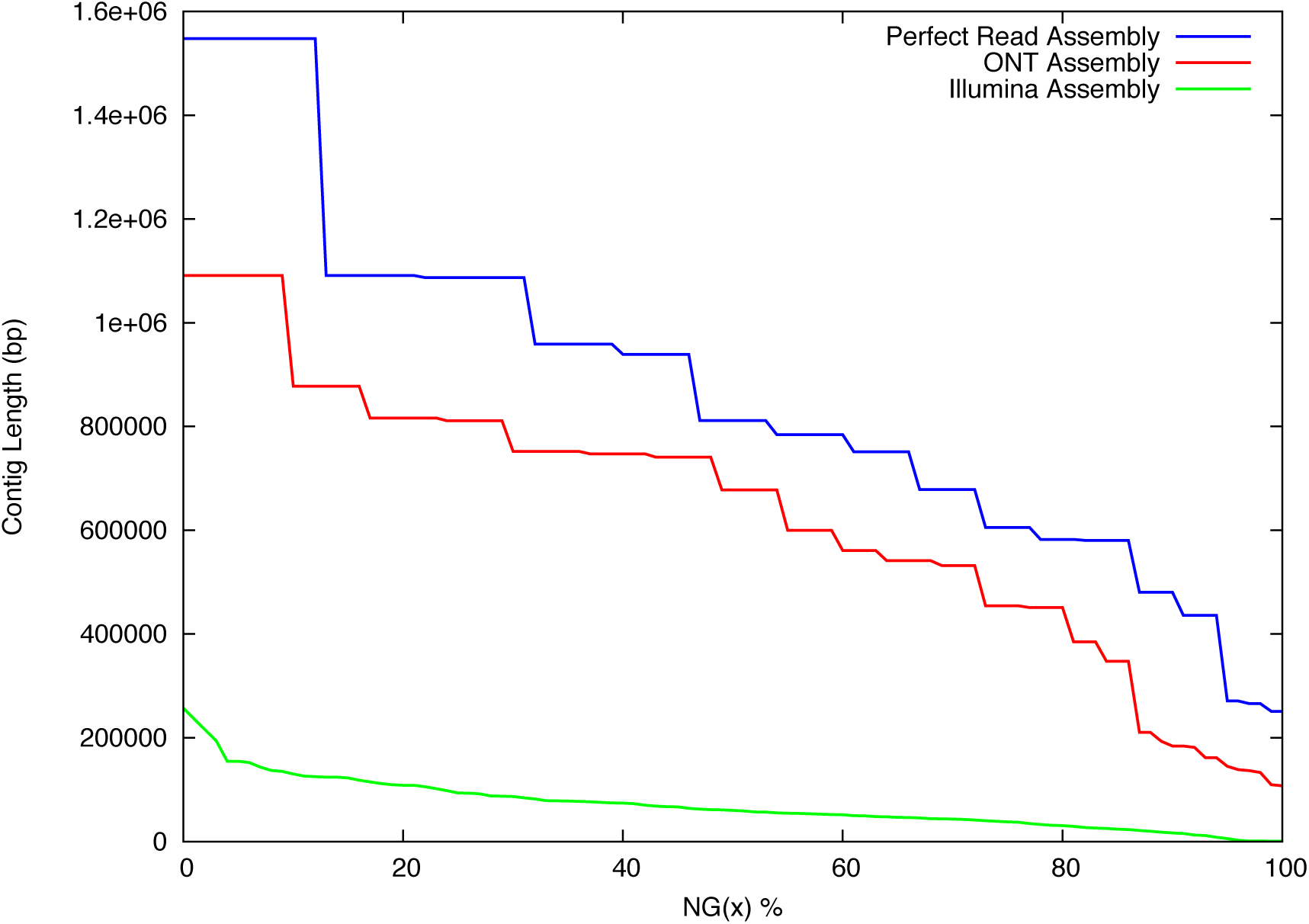
NG-graph Comparison of corrected Oxford Nanopore assembly and an Illumina assembly. NG-graph of a simulated perfect read assembly, the corrected Oxford Nanopore assembly and an Illumina-only assembly. The curve extends the common N50 metric to trace the contig size such that the top X% of the genome is assembled into contigs this size or larger. The Oxford Nanopore assembly is substantially more contiguous across the entire size spectrum and is far closer to the perfect read assembly than the Illumina-only assembly. Notably, the N50 contig length for the Oxford Nanopore-based assembly is 678kbp compared to about 60kbp for the Illumina assembly and is quite close to the 811kbp perfect read assembly N50.

To evaluate the effectiveness of the hybrid error correction and assembly algorithm, we also computed a “reference-based” assembly of the Nanopore reads by extracting sequences from the reference as “perfect reads” where the Nanopore reads aligned. Interestingly, assembling these “perfect reads” lead to nearly the same results: the contig N50 was at best 811kbp for the reference assembly compared to 678kbp for the Nanocorr-corrected reads. This highlights that the remaining contig breaks in the Nanocorr assembly were due to the sequence composition and repeat structure of the genome and to a much lesser degree the small amount of residual error after correction **(Supplementary Note 8)**. Finally, to evaluate the minimum amount of raw coverage needed to achieve these results, we computed 46 separate assemblies using the top N most productive flowcells. We find that the best result was achieved by using the data from just the top 3 flowcells, representing ∼30x raw coverage of the genome **(Supplementary Note 10)**.

We sought to observe the differences in biological insights that could be obtained by the analysis of genome assemblies with different degrees of underlying contiguity. Aligning the Illumina and Oxford Nanopore/Illumina hybrid assemblies against the reference yeast genome allowed us to evaluate how well the two assemblies represented the various classes of annotated genomic features. While both the Illumina-only and nanopore-based assemblies could correctly assemble short genomic features, the nanopore-based assembly was able to substantially outperform the Illumina-only assembly of the longest genomic features (**Figure 4**). In particular, rRNAs (averaging 1393bp), gene cassettes (averaging 2951bp), telomeres (averaging 4396bp), and transposable elements (averaging 3201bp) were substantially better represented in the nanopore assembly, and nearly completely absent from in Illumina-only assembly. Only the very longest repeats in the genome, such as the 20kbp telomeric repeats, remain unresolved in the Oxford Nanopore assembly and become fragmented in both assemblies as well as the reference-based assembly. The MiSeq assembly slightly outperforms for “binding site” features, although these are binding sites within the telomeric repeats that were not well assembled by either technology.

**Figure 4.**
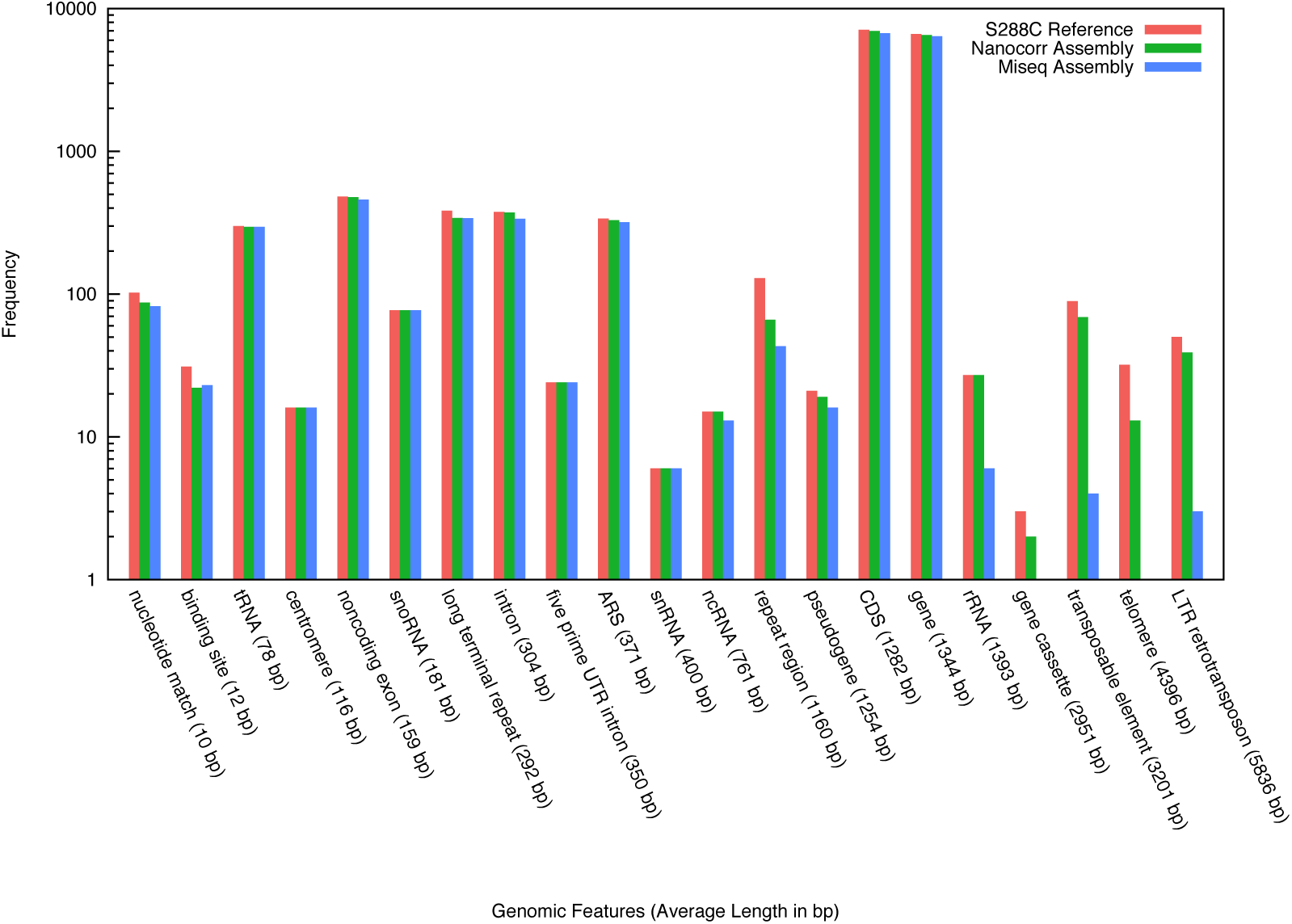
Genomic features assembly by Oxford Nanopore and Illumina sequencing. Quantification of different annotated genomic features assembled completely by the Nanopore and Illumina/MiSeq only assembly relative to the complete S288C reference annotation. The Nanopore-based assembly produces an assembly with many more of the longer and repetitive features assembled compared to the Illumina/MiSeq-only assembly.

### E. coli K12 Error Correction and Assembly

In order to validate the utility of this workflow, we also error corrected and *de novo* assembled the Oxford Nanopore reads generated by Nicholas Loman et al.^15^ of *E. coli* K12 using the same approach (**Supplemental Note 7**). In this experiment, a total of 145x Oxford Nanopore read coverage of the genome was error corrected with the Nanocorr pipeline using 30x Illumina MiSeq coverage to improve the average identity to over 99%. This time, only reads greater than 7kb in length, representing ∼28x coverage of the genome, were used in the assembly. The final result was an essentially perfect single 4.6Mbp chromosome length contig with >99.99% identity. In contrast, the Illumina-only assembly produced an assembly with hundreds of contigs and a contig N50 size of only 176kbp.

## Discussion

The results of this study indicate the Oxford Nanopore sequence data currently have substantial errors (∼5% to 40% error) and a high proportion of reads that completely fail to align (∼50%). This is likely due to the challenges of the signal processing the ionic current measurements^16^ as well as the challenges inherent in any type of single molecule sequencing. Oxford Nanopore has indicated that the pores are more than a single base in height so that the ionic signal measurements are not of individual nucleotides but of approximately 5 nucleotides at a time. Consequently the base calling must individually recognize at least 4^5^=1024 possible states of ionic current for each possible *5-mer*. We also observed the potential for some bias in the signal processing and basecaller, particularly for homopolymers (**Supplemental Figure S4**). Despite the limitations of this early phase device, there has been notable improvement over the course of this program, and well performing flowcells of the current iteration (R7.3 at the time this publication was written) can generate upwards of 400Mb on a single run. Continuing this improvement of yields with future generations of the technology would obviously add considerably to the utility of the system.

While short read sequencers in general have lower error rates and, to date, have become the standard approach of genomics, short reads are not sufficient to generate long continuous assemblies of complex genomes. To this day the reference human genome remains incomplete as do the reference genomes for most higher species, especially plants. Long reads are necessary to span repetitive elements and other complex sequences to generate high quality, highly contiguous assemblies. Currently there are limited methods for generating adequately long reads. Synthetic long reads can be generated on existing short read platforms using barcoding approaches such as those employed by Illumina’s TruSeq Synthetic Long-Read approach (formerly Moleculo) and the new 10X Genomics platform, however these approaches still rely on the existing short read infrastructure. Alternatively, true long reads can be generated by the Pacific Bioscience System and now the Oxford Nanopore MinION.

Improving the contiguity of a genome assembly enables more detailed study of its biological content and function in every aspect. Genes will more often be correctly assembled along with their flanking sequences, enabling deeper study of regulatory elements. Longer reads will also resolve more repetitive sequences as well, especially transposable elements, high copy genes, segmental duplications, and centromeric/telomeric repeats that are difficult to assemble with short reads. Finally, high-quality assemblies are also essential to study high-level genome structures such as the evolution and synteny of entire chromosomes across species. Even in genome resequencing, short reads can be problematic, with some (perhaps many) structural variants unresolved, obscuring the true gene content of a member of a species or obscuring clinically relevant structural variants in an affected individual^17^.

Modern genome assemblers are not equipped to natively handle reads with error rates above a few percent. Consequently, before the Oxford Nanopore reads can be used for *de novo* assembly they must first be error corrected. These general strategies are helpful for other single molecule, long read sequences such as those from Pacific Biosciences, although existing algorithms were not capable of resolving the Oxford Nanopore errors^17,18^. We successfully developed a new hybrid error correction approach that can improve the average per base identity of the Oxford Nanopore reads from 65% across all flowcell iterations to greater than 97% and generates nearly perfect or extremely high quality assemblies given sufficient coverage and read lengths. Using the error corrected data, we were able to fully reconstruct an entire microbial genome and produce an extremely high quality assembly of yeast that had many important genomic features that were almost entirely lost in the Illumina-only assembly. This work has demonstrated how single molecule, long read data generated by the Oxford MinION can be successfully used to compliment short read data to create highly contiguous genome assemblies, paving the way for essentially any lab to create perfect or high quality reference sequences for their microbial or small eukaryotic projects using a handheld long read sequencer.

## Methods

### Yeast growth

An aliquot of yeast strain W303 was obtained from Dr. Gholson Lyon (CSHL). Four ml cultures in 15 mL falcon tubes of yeast were grown in YPD overnight in at 32°C to ∼1×10^8^ cells. The cells were purified using the Gentra Puregene Yeast/Bacteria kit (Qiagen, Valencia CA). DNA was stored at - 20°C for no more than 7 days prior to use.

### Library preparation

#### Oxford Nanopore

Purified DNA was sheared to 10kb or 20kb fragments using a Covaris g-tube (Covaris, Woburn MA). Four ug of Purified DNA in 150 ul of DI water was loaded into a g-tube and spun at 6000 RPM Eppendorff 5424 for 120 sec (10kb) or 4200 RPM for 120 sec (20kb). All DNA was further purified by adding 0.4X AMPure beads. A twisted kimwipe was used to remove all visible traces of ethanol from the walls of the tube. The beads were allowed to air dry and DNA was eluted into 30ul of DI water.

#### R6.0 and R7.0 preparation

The DNA concentration was measured with a Qubit fluorometer and an aliquot was diluted up to 80 ul. Five ul of CS DNA (Oxford Nanopore, Oxford UK) was added and the DNA was end-repaired using the NEBNext End Repair Module (NEB, Ipswich MA). The DNA was purified with AMPure beads and eluted in 25.2 ul of DI water. DNA A-tailing was performed with the NEBNext dA-Tailing module (NEB, Ipswich MA).

Blunt/TA ligase (NEB, Ipswich MA) was added to the A-tailed library along with 10 ul of the adapter mix (ONT, Oxford UK) and 10 ul of HP adapter (ONT, Oxford UK). The reaction was allowed to incubate at 25°C for 15 minutes. The DNA was purified with 0.4X of AMPure beads. After removal of supernatant, the beads were washed 1X with 150 ul Wash Buffer (ONT, Oxford UK). After supernatant was removed the beads were briefly spun down and then re-pelleted and the remaining supernatant was removed. A twisted kimwipe was used to remove all traces of wash buffer from the wall of the tube. The DNA was resuspended in 25 ul of Elution Buffer (ONT, Oxford UK).

The DNA was quantified using a qubit to estimate the total ng of genomic+CS DNA in the final library. Ten ul of tether (ONT, Oxford UK) was added to the ligated library and allowed to incubate at room temperature for 10 minutes. Fifteen ul of HP motor was then added and allowed to incubate for 30 minutes or overnight.

Between 5 and 250 ng of the pre-sequencing library was diluted to 146 ul in EP Buffer (ONT, Oxford UK) and 4 ul of Fuel Mix (ONT, Oxford UK) was added to the sequencing mix. The library was immediately loaded on to a flow cell.

#### R7.3 preparation

The DNA concentration was measured with a Qubit fluorometer and an aliquot was diluted up to 80 ul. The DNA was end-repaired using the NEBNext End Repair Module (NEB, Ipswich MA). The DNA was purified with AMPure beads and eluted in 25.2 ul of DI water. DNA A-tailing was performed with the NEBNext dA-Tailing module (NEB, Ipswich MA).

Blunt/TA ligase (NEB, Ipswich MA) was added to the A-tailed library along with 10 ul of the adapter mix (ONT, Oxford UK) and 2 ul of HP adapter (ONT, Oxford UK). The reaction was allowed to incubate at 25°C for 15 minutes. The DNA was purified with 10ul of his-tag Dynabeads (Life Technologies, Norwalk CT) suspended in 100ul of 2X Wash Buffer (ONT, Oxford, UK). After removal of supernatant, the beads were washed 2X with 250 ul of 1X Wash Buffer (ONT, Oxford UK). After supernatant was removed the beads were briefly spun down and then re-pelleted and the remaining supernatant was removed. A twisted kimwipe was used to remove all traces of wash buffer from the wall of the tube. The DNA was resuspended in 25 ul of Elution Buffer (ONT, Oxford UK). The DNA was quantified using a qubit to estimate the total ng of genomic DNA in the final library.

Between 5 and 250 ng of the pre-sequencing library was diluted to 146 ul in EP Buffer (ONT, Oxford UK) and 4 ul of Fuel Mix (ONT, Oxford UK) was added and the the sequencing mix. The library was immediately loaded on to a flow cell.

Libraries were sequenced using the MinION device for between 48 and 72 hours. Whenever possible, DNA was handled with a wide bore, low bind pipette tip. Mixing of DNA with reagents was done by flicking or preferably pipetting with a wide bore tip. All tube used were Protein LoBind (Eppendof, Hamburg Germany) All material loaded onto a flow cell was loaded using a 1000 up pipette. Deviations from this protocol for each flow cell can be found in supplemental methods.

#### MiSeq

One ug of yeast DNA purified using the Gentra Puregene Yeast/Bacteria kit (Qiagen, Valencia CA) was prepared using a TruSeq PCR free kit (Illumina). The insert size was 350 with a paired end 250 run.

### Flowcell disposition

Flowcell were received on ice and immediately stored at 4°C. Ideally within 3 days each flow cell was QC’d with the minKnow software and the number of available pores was recorded. The flowcells with 400 available pores or more were generally considered “good” and used first. Immediately prior to library loading the flowcell was removed from the 20°C refrigerator and flushed with 150ul of EP Buffer (ONT). The flowcell was allowed to incubate at room temperature for 10 minutes followed by a second EP flush and incubation.

For flowcells that were washed prior to the addition of additional library; the flow cells were washed with 150 ul of Solution A (ONT) followed by a 10 minute room temperature incubation. One-hundred and fifty ul for solution B (ONT) was then added and the flowcells were stored at 4°C until use. Prior to use the washed flowcells were flushed with EP Buffer (ONT) as previously described.

### Read Alignment and Error Characteristics

Yield over time data extraction, individual flow cell statistics calculation, fasta/fastq generation were all performed using poretools^19^. Plots were generated using R (ggplot2) and gnuplot. Overall accuracy was calculated by aligning the raw Oxford Nanopore reads to the W303 pacbio assembly using Blast version 2.2.30+ with the following parameters:

-reward 5 -penalty -4 -gapopen 8 -gapextend 6 -task blastn -dust no -evalue 1e-10 High Scoring Segment Pairs were filtered using the LIS algorithm and a scoring function that penalizes overlaps while maximizing alignment lengths and accuracy. Overall accuracy was calculated by averaging the percent identity of all of the filtered HSP’s derived from all of the reads. Error rate over the read length was calculated by taking the HSP’s from a sampling of 1000 random reads in the dataset with read lengths between 9kb and 10kb. The identity was calculated for 100bp sliding windows over the length of the alignment and averaged over all of the alignments.

### Read Correction and Assembly

Raw reads were extracted from the h5 files generated by the basecaller. Only independent reads, one per molecule, were corrected for assembly. Because a channel can produce three reads of the same molecule, reads were chosen in order of their expected accuracy: 2D or the 1D template to represent each DNA fragment. As part of the Nanocorr algorithm, 30x coverage of 300bp paired end MiSeq data was then aligned to the nanopore reads using blastn with the following parameters:

-reward 5 -penalty -4 -gapopen 8 -gapextend 6 -task blastn -dust no -evalue 1e-10 Nanopore reads to which no MiSeq reads aligned were excluded from the process. The Nanocorr algorithm then filters the alignments by first removing those contained within a larger alignment and then an LIS Dynamic Programming algorithm was applied using a scoring scheme to minimize the overlaps in the alignments. The filtered set of alignments was then used to build a consensus using ‘pbdagcon’^13^ (https://github.com/PacificBiosciences/pbdagcon.git). The software and documentation for the error correction software are available open source at https://github.com/jgurtowski/nanocorr.

The error correct nanopore reads were then assembled using Celera Assembler version 8.2beta (http://wgs-assembler.sourceforge.net/). Redundant contigs, representing individual nanopore reads with higher rates of residual errors were then identified using ‘blastclust’ (which is part of the blast executable package found at http://blast.ncbi.nlm.nih.gov/Blast.cgi?PAGE_TYPE=BlastDocs&DOC_TYPE=Download). This algorithm identifies sequences that align to the interior of another longer sequence, using the parameters:

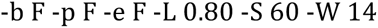

Finally, the non-redundant contigs were then ‘polished’ using the Pilon algorithm that revises the consensus sequence using the alignments of the MiSeq reads to the newly assembled contigs.

Alignments and dotplots were generated using ‘nucmer’ and ‘mummerplot’ from the MUMmer version 3.23 package^20^.

### Feature Quantification

Each assembly was aligned to the S288C reference genome using *nucmer* from the MUMmer version 3.23 package. Alignments were filtered using the command *delta-filter -1*, also from the MUMmer 3.23 package to find the best non-redundant set of contigs. The non-redundant set of alignments was intersected with the feature coordinates from the S288C annotation obtained from the Saccharomyces Genome Database using BEDTools^21^ command *intersectBed* with the parameters : -u –wa –f 1.0. The features that were fully contained in an alignment were included in the tally seen in Figure 4.

### Data Access

All data and software used in the study are available open source at: http://schatzlab.cshl.edu/data/nanocorr/

## Acknowledgments

This project was supported in part by National Science Foundation award DBI-1350041 and National Institutes of Health award R01-HG006677 to MCS. We would like to express our thanks to Oxford Nanopore for affording us the opportunity to participate in the MinION early Access program (MAP). In particular we would like to thank Clive Brown, James Brayer and all the members of the technical support staff for their support and assistance during this research. Finally, we would like to thank all the members of the MAP community for their on-going insight and dedication into the novel device.

## Author contribution

SG Performed data analysis, library preparation, managed flow cells and is the MAP lead. JG Performed data analysis, developed Nanocorr and library preparation. SE performed library preparation, PD performed library preparation. MCS assisted in data analysis and in the overall design of the project. WRM developed the overall design of the study and assisted with library preparation. SG, JG, MCS and WRM wrote the manuscript. All authors reviewed and approve the final manuscript.

